# Berberine: A dual anti-HIV and anti- cervical cancer compound

**DOI:** 10.1101/2024.11.18.624212

**Authors:** Wasifa Naushad, Bryson C. Okeoma, Humayun K Islam, Zhen Zhen Wang, Nan Yang, XiuMin Li, Chioma M. Okeoma

**Author notes:** These authors contributed equally to this work (Wasifa Naushad, Bryson C. Okeoma). (Wasifa Naushad), (Bryson C. Okeoma), (Humayun K Islam), (Zhen Zhen Wang), (Nan Yang), (XiuMin Li), (Chioma M. Okeoma).

## Abstract

We report the effects of berberine (BBR), a benzylisoquinoline alkaloid small molecule on inhibition of HIV infection of cervical cancer cells. We used HeLa cell-derived TZM-bl cell line as a model of cervical cancer and HIV infection. BBR significantly inhibits viral and cancer processes, including expression of cell-associated HIV RNA, secretion of HIV reverse transcriptase, and HIV Tat-mediated LTR promoter transactivation. BBR significantly inhibits HIV-induced cancer cell viability and cell clustering. Besides its ability to inhibit HIV-induced cancer cell viability, BBR inhibits migration and matrix invasion of cervical cancer cells that are infected with HIV or treated with HIV Tat protein. The ability of BBR to inhibit cell clustering, collective cell migration, and cell invasion may have an effect in controlling progression of cervical cancer and HIV since collective migration and invasion are strategies for local tissue infiltration, as well as metastatic invasion in epithelial cancers. Interestingly, molecular docking and dynamic stimulation show that BBR binds HIV_IIIB_ Tat amino acid residues through non-covalent interactions that occurs at multiple sites including LYS71, providing mechanistic insights into BBR regulation HIV infection. The results of the present study suggest that BBR has the potential to inhibit HIV infection and comorbid cervical cancer progression.

**Graphical abstract:** 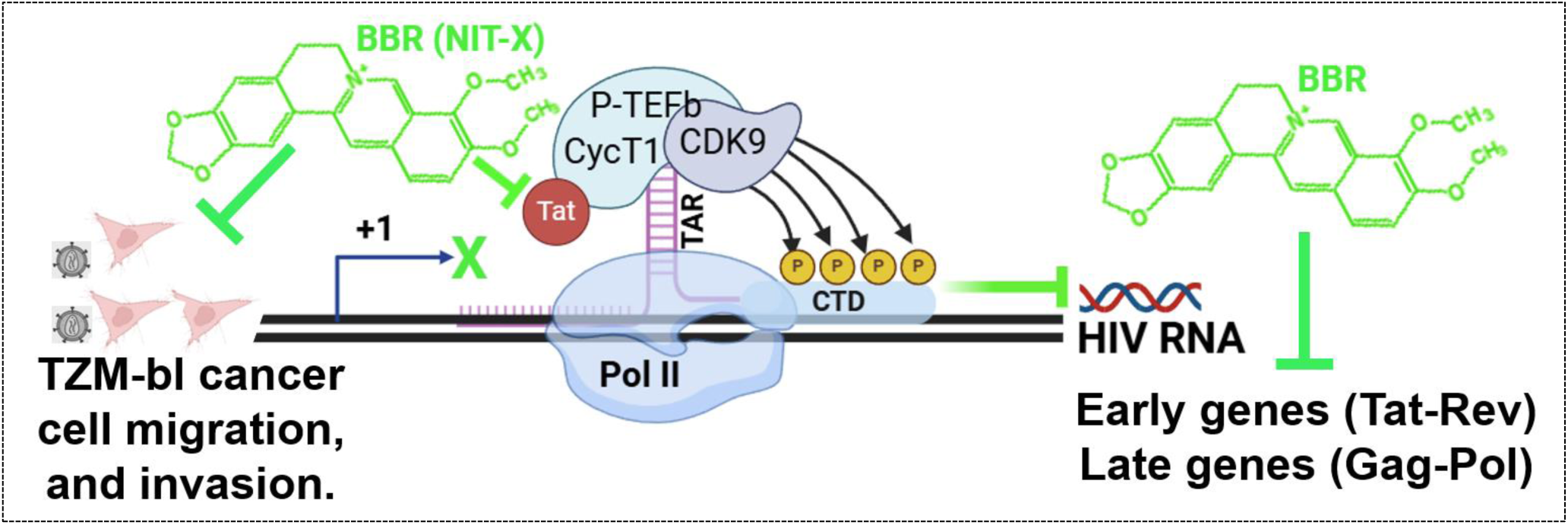

Berberine (BBR), a benzylisoquinoline alkaloid small molecule, inhibits HIV infection of cervical cancer cells, suppresses HIV-induced migration and invasion of the cancer cells. Mechanistically, BBR interferes with Tat-mediated HIV LTR promoter transactivation and expression of different HIV RNA species, expression of early genes (multiply spliced Tat-Rev) and late genes (unspliced Gag-Pol).

## 1. INTRODUCTION

Cervical cancer is amongst the acquired immunodeficiency syndrome (AIDS)-defining cancers [1], as HIV infected women have a higher risk of developing cervical cancer compared with HIV uninfected women [2, 3]. Human papillomavirus (HPV) is a small nonenveloped double-stranded DNA virus that is associated with almost 100% of all cervical cancers. HPV is classified into mucosal and cutaneous HPVs and designated as high risk (HR) or low risk (LR) HPV according to their propensity for malignancy. Several epidemiologic studies have conclusively demonstrated that infection with HR-HPV is the primary risk factor for the development of cervical cancer and its precursors [4]. The oncogenic potential of HR-HPV is highly linked to the activities of E6 and E7 oncogenes that abrogate the functions of p53 and pRB, respectively [5, 6]. However, HR-HPV infection alone is not enough to cause cancer, indicating that there are additional components that play roles in HPV-induced cancer, such as the oncogenic activities of HIV and HIV proteins, such as Tat [7–11].

HIV Tat protein is an early viral protein necessary for virus replication. Tat is secreted by infected cells and is taken up by uninfected cells, including epithelial cells [12]. Tat has been implicated in increased cell migration and invasion [13], cell proliferation and differentiation [14], as well as interaction with cell cycle proteins [7, 15]. Tat directly interacts with the tumor suppressors -retinoblastoma protein (pRb) RB2/p130 [16, 17] and E2F4 [18]. Additional oncogenic activities of Tat include promotion of cervical cancer cell survival and proliferation via upregulating the expression of proliferation marker cyclin A and decreasing levels of cell cycle inhibitors [7]. Furthermore, the oncogenic activities of Tat may depend on its ability to phosphorylate the focal adhesion kinases [19, 20], promote cellular adhesion [21, 22], cellular migration [21, 23], and induction of the synthesis of matrix metalloproteases (MMPs) [24, 25]. Thus, Tat has pleiotropic effects on cellular functions and these diverse effects of Tat may be necessary in reprograming cervical cancer cells.

The advent of antiretroviral therapy (ART) has increased the lifespan of people with HIV (PWH) [26]. However, it is not clear whether ART reduces the incidence of cervical cancer or not. In PWH comorbid with HPV, ART was shown to play significant role in clearing HR-HPV in anal mucosa of men who have sex with men (MSM) [27]. In some women, ART was associated with significant reduction in the risk of cervical lesion progression in PLW [28], while in others, the incidence of cervical cancer risk is still significantly higher in PWH compared to the general population [29]. Since about 5.2% of all cervical cancers in women are attributable to HPV-induced intraepithelial lesions, and HIV exacerbates the disease progression, HIV infection constitutes a significant disease and economic burden worldwide, especially in low-income countries with limited access to ART.

Despite the effectiveness of ART in suppressing viremia, ART is not a cure since it does not stop basal viral transcription and production of viral proteins. Additionally, the efficacy of ART is often limited by drug toxicity, poor adherence to therapy, and possible drug resistance. In spite of these shortcomings, access to ART remains a major hurdle to its use by PWH in developing countries. There is evidence that the use of TCM as complementary and alternative medicine (CAM), reduces the side effects of ART and improves immunity, clinical symptoms, and quality of life of PWH [30, 31]. However, there is insufficient evidence for the efficacy of TCM in regulating HIV promoter transactivation and expression of CA-HIV RNA, especially in people with cervical cancer. Hence, a rigorously designed cell-based assay is needed to assess the effectiveness of TCM on HIV promoter transactivation and expression of CA-HIV RNA in cervical cancer cells.

In this study, HeLa cell-derived TZM-bl reporter cell line was used as a model to study the effects of BBR, a benzylisoquinoline alkaloid small molecule compound isolated from medicinal plants and known to have anti-diabetic, anti-inflammatory, and anti-viral activities [32–34] on HIV infection of cervical cancer cells. TZM-bl cell line is a reporter cell line (JC57BL-13) that was engineered from the cervical cancer cell line (HeLa cells) to express HIV receptor (CD4) and coreceptors (CXCR4 and CCR5) [35], and the cells were also engineered with two lentiviral vectors coding for the reporter genes—firefly luciferase and *E. coli* β-galactosidase that are under the control of the HIV-1 LTR [35, 36]. As a result, TZM-bl cells are sensitive to infection with different isolates of HIV-1 [37–39]. We assessed the effects of BBR on HIV LTR promoter transactivation, expression of CA-HIV RNA, and secretion of HIV reverse transcriptase (RT) as indicators of HIV infection, while cell viability, migration, and invasion were used as hallmarks of cervical cancer cells.

### Impact

This study demonstrates that BBR has the potential to inhibit coincident HIV promoter activation, expression of CA-HIV RNA, viability, migration, and invasion of HIV infected TZM-bl cervical cancer cells. BRB also inhibits HIV Tat protein mediated transactivation of HIV LTR promoter, suppress the expression of Tat mRNA, and BBR interacts with HIV subtypes B and C, and binds HIV_IIIB_ subtype B Tat amino acid residues through non-covalent interaction that occurs at multiple sites including LYS71. If validated, the use of BBR may provide some relief in PWH with limited access to ART who are comorbid with cervical cancer, especially people in the African continent and other parts of the world with similar predicaments. A safe application of this potential alternative therapy could help manage disease burden in PWH and cervical cancer, which are in agreement with 2030 Sustainable Development Goals to reduce mortality from non-communicable diseases through prevention and treatment.

## 2. MATERIALS AND METHODS

### 2.1 High-performance liquid chromatography analysis of compounds

Berberine (BBR) was obtained from Xi’an SaiYang biotechnology, LLC (Xi’an, P.R. China). 7,4′-dihydroxyflavone (DHF) was obtained from Biosynth (Gardner, MA). Isoliquiritigenin (ILG) was purchased from Xi’an QinMing Biotech Co., Ltd (Xi’an, P.R. China). HPLC analysis of these 3 compounds was carried out on a Waters 2690 HPLC system coupled with a 2996 PDA detector (Waters, Milford, MA). All three compound were dissolved in methanol at the concentration of 100 µg/mL, and 10 µL of the sample solution was injected into the HPLC system. 0.1% formic acid solution was used as mobile phase A and HPLC grade acetonitrile (Fisher Scientific, NJ) was used as mobile phase B. A ZORBAX SB-C18 (4.6 × 150 mm, 5 µm) column (Agilent, Santa Clara, CA) was used for the separation analysis under the mixed mobile phase at a flow rate of 1 mL/min. The gradient elution process was as follows: from 10% mobile phase B to 100% in 10 minutes and maintain at 100% mobile phase B for 3 minutes. The chromatographic data was acquired and processed using Waters Empower software.

### 2.2 Cells, HIV, and Tat protein

TZM-bl cell line was obtained through the Biodefense and Emerging Infections Research Resources Repository ([BEI-RRP] https://www.beiresources.org/, formerly NIH AIDS Reagent program) and were maintained in DMEM (Gibco-BRL/Life Technologies) containing 10 % FBS (Gibco), 100 U/ml penicillin, 100 μg/ml streptomycin, sodium pyruvate and 0.3 mg/ml L-glutamine (Invitrogen, Molecular Probes) as previously described [38]. HIV-1 Infected U937 Cells (U1), ARP-165 (BEI-RRP) contributed by Folks et al., 1989 [40] were grown in RPMI (Gibco-BRL/Life Technologies) containing 10 % FBS (Gibco), 100 U/ml penicillin, 100 μg/ml streptomycin, sodium pyruvate, and 0.3 mg/ml l-glutamine (Invitrogen, Molecular Probes) as previously described by [41]. HIV_U1_ was purified from supernatants from U1 cells. HIV-1Tat protein (ARP-2222) is a full length, biologically active recombinant protein derived from HIV-1_IIIB_ Tat protein obtained through BEI-RRP. HIV-1_IIIB_ Tat protein was produced in an *E. coli* expression system, purified by affinity chromatography on heparin sepharose, followed by reverse phase chromatography, and contributed by DAIDS/NIAID.

### 2.3 Infection of TZM-bl cervical cancer cells with HIV and treatment with TCM compounds

Equivalent numbers (10,000) of TZM-bl cells/well were plated in 100 µL of DMEM media in a 96 well plate. Cells were allowed to adhere to the plate for 5 hours and then infected with HIV_U1_ containing 8 RT units/µL in 10 µL. 24 hours later or as indicated in the figure legends, cells were treated with different concentrations (20, 40, 80 and 100 µg/mL) of TCM compounds. Vehicle (DMSO) was used as control in place of TCM compounds. 24 hours after treatment with TCM compounds or as indicated in the figure legends, cells were used for various assays, including analysis of cell viability, HIV promoter activity, expression of CA-HIV RNA species (Tat-Rev, Gag-Pol RNA).

### 2.4 Production of HIV and assessment of HIV reverse transcriptase (RT) assay

2.5 × 10^6^ HIV infected U1 cells were plated in 500 µL of media in a 24 well plate overnight. 24 hours later, culture supernatants were collected and clarified by centrifugation. The virus produced was quantified by the standard RT assay. The virus was aliquoted and stored at −80 °C at 8 RT units/µL in 10 - 15 µL.

### 2.5 LTR promoter activation assay

Vehicle (DMSO), HIV_U1_, Tat 0.4 µg, BBR 40 μg, or 100 µg, were added to TZM-bl indicator cells. Infection or treatment with Tat was allowed to proceed for 24 hours before initiation of treatment with TCM compounds for another 24 hours. Cells were then assessed for HIV promoter LTR transactivation by measuring luciferase reporter activity as previously described [39]. Briefly, cells to be analyzed were removed from the 37°C incubator and equilibrated to ambient temperature (22 to 25°C) on the bench for 5 to 15 minutes. Using a manual multichannel pipette, a 100 µL volume of Bio-Glo™ Reagent equal to the volume of cells only or cells/test sample mixtures to each assay well. Assay plates were incubated at ambient temperature for 5 to 15 minutes. Luminescence was measured using BioTek Synergy H1 Microplate Reader.

### 2.6 BBR cellular uptake assay

To assess the ability of cervical cell to internalize BBR, we used the fluorescent properties BBR in our cellular internalization assay [42]. Briefly, equivalent numbers (5000 cells/well) of cells were seeded in each well (3 wells per 3 independent experiments) of a 96-well plate. Cells were treated with BBR 100 μg /mL or equivalent volume of DMSO (negative control) and incubated for 1 hour. Images of cells with internalized fluorescent BBR were obtained after 1 hour using BioTek Lionheart FX automated microscope at 10x. Fluorescent intensity was computed using Gen5 software and plotted using Prism GraphPad version 10.

### 2.7 Assessment of cell viability on monolayer

TZM-bl cells were seeded at 5000 cells/well in a 96-well plate and cultured overnight at 37 °C, 5% CO_2_. 24 hours later, cells were treated with 50 µl of DMSO used as vehicle (control) or different concentrations (40 µg or 100 µg) of compounds in 50 µl of media. The cells were then cultured for 24 hours or 48 hours, after which each well was treated with 20 μl of MTT (3-[4,5-dimethylthiazol-2-yl]-2,5 diphenyl tetrazolium bromide) reagent. Following the addition of MTT, the plate was incubated at 37°C for 3 hours. After 3 hours, 100 μl of MTT solvent (90% Isopropanol, 10% Triton X-and 0.4% 1M HCl) was added to each well and the plate was covered from light while slowly shaking for 30 minutes. The absorbance of solubilized MTT reagent was measured using a spectrophotometer at 590 as we previously described [42, 43].

### 2.8 Cell Clustering

The wells of a 96-well tissue-culture plate were seeded with TZM-bl cells (5000 cells/well) and cultured overnight at 37 °C, 5% CO_2_. 24 hours later, cells were left uninfected or infected with HIV (8 RT units/µL) or treated with 40 µg of BBR in 50 µl of media or 50 µl of DMSO alone, used as vehicle (control). The following day, cells were imaged for visual analysis of cell clustering using BioTek Lionheart FX automated microscope at 10x.

### 2.9 Isolation of RNA and assesment of gene expression by quantitative PCR

Total RNA was isolated from cells using Zymogen Quick RNA isolation kit, per manufacturer’s protocol. RNA was eluted and the eluate was measured using a NanoDrop 1000. 1 µg of total RNA was used for cDNA synthesis. 500 µg of cDNA was used for analysis of gene expression by real-time quantitative PCR (RT-qPCR). RT-qPCR was permormed using a 7500 FAST machine and Power Track SYBR Green master mix as previously described using the primers listed on **Table 1**.

**Table 1.**
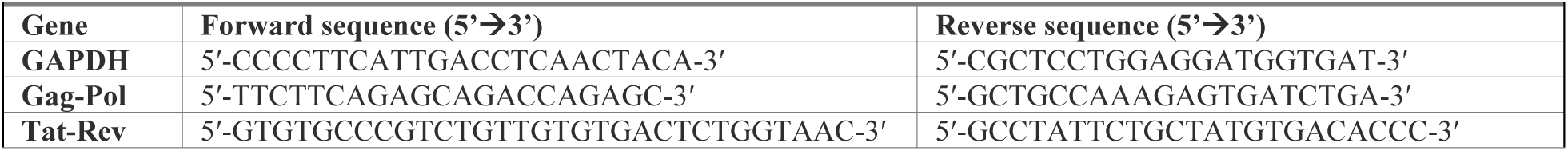
Primer sequences used in this study.

### 2.10 Cell migration by wound healing assay

To perform the cell migration assay, we used our previoulsy published protocol [43]. 10,000 TZM-bl cells were plated in a 12 well plate. For the assessment of the effects of BBR on HIV-induced migration of cervical cancer cells, TZM-bl cells were left uninfected or infected with HIV_U1_ for 6 hours in complete media (DMEM supplemented with 10% FBS). For assessment of the effects of BBR on migration of HIV Tat induced cervical cell migration, TZM-bl cells were transfected with pCMV-TAT plasmid (2 µg) with Lipofectamine™ 3000 Transfection Reagent (Invitrogen Cat# L3000001), and plates were incubated at 37 °C with 5% CO_2_ for 24 hours. After 6 hours of infection or 24 hours of transfection, media was removed, and cells were cultured in fresh complete media containing BBR (40 µg/mL or 100 µg/mL) or 0.1X DMSO. The scratch (wound) was created using 10 µL pipette tip across the middle of the plate. Cells were incubated at 37 °C with 5% CO_2_ for 3 hours. The wound areas were imaged at different time points (0, 6, 18 and 24 hours) to quantify the rate of wound closure, which signifies cell migration. Images were taken with Lionheart-FX microscope with 10 X objective. Images were processed using Gen5 software.

### 2.11 Preparation of transwell inserts for cell invasion

Transwell inserts (Corning Cat# 3415, with 3.0 µm Pore size) were coated with 3mg/mL of Matrigel (Corning, Cat#356234) in 100 µL of serum free media. The inserts were incubated at 37 °C for 3 hours prior to use.

### 2.12 Transwell invasion assay

10,000 TZM-bl cells/well were transfected with pCMV-TAT plasmid (2 µg) with Lipofectamine™ 3000 Transfection Reagent (Invitrogen Cat# L3000001), and plates were incubated at 37 °C with 5% CO_2_ for 24 hours. After 24 hours, media was removed, cells were washed, and BBR (40 µg/mL and 100 µg/mL) or 0.1X DMSO (vehicle) added. Cells were cultured for 24 hours after which the complete media was removed and serum free media added to starve the cells. After 3 hours in serum free media, cells were trypsinized and washed. 100,000 cells were added to the apical side of Matrigel-coated Transwell inserts in serum free media. Inserts were then placed in a 24 well plate containing DMEM supplemented with 10% FBS with 10µg/mL of fibronectin (basal chamber). Cells were incubated at 37 °C with 5% CO_2_ for 3 hours. After 3 hours, 100 µL of complete media containing 10% FBS was added to the apical chamber. Cells were incubated at 37 °C with 5% CO_2_ for 24 hours. After 24 hours of incubation, inserts were removed and washed to remove loose cells. Inserts were Giemsa stained to visualize membrane-bound cells. Images were taken with Lionheart-FX microscope with 10 X objective. Images were processed using Gen5 software.

### 2.13 The sources of the sequences of HIV Tat used for Docking and molecular dynamics simulation

The HIV-1 subtype B (Isolate MN, ACCESSION P05905, https://www.uniprot.org/uniprot/P05905) and subtype C Tat (Isolate 92BR025, ACCESSION O12161, https://www.uniprot.org/uniprot/O12161) sequences used for docking and molecular dynamics analyses were retrieved from the Universal Protein Databases (UniProt). These Tat subtypes were used because it has been shown that variations in their sequences maybe related to differential effects of HIV on neuropathogenesis [44].

### 2.14 Docking and molecular dynamics

The HIV subtype B protein (TatB) sequences were obtained from the Universal Protein Databases (UniProt). The Swiss-Model server was applied to model the three-dimensional structure of TatB. The structure of berberine is strived from Pubchem database. The docking between berberine and TatB were carried out utilizing Autodock Vina. The low-energy configurations of berberine bound to TatB were selected for the further molecular dynamic simulation. The molecular dynamics simulations were executed utilizing the Groningen Machine for Chemical Simulations (GROMACS) software, incorporating the amber99sb-ildn force field, the GAFF (Generalized Amber Force Field) for berberine, and the TIP3P water model for solvent representation. Energy minimization with steepest descents, NVT, and NPT ensembles have been engaged in order for optimizing the system. V-rescale temperature-coupling method was adopted for NVT. Parrinello–Rahman pressure coupling was applied with constant coupling for NPT. Electrostatic forces were calculated by the Particle Mesh Ewald method. The simulation of the TatB-berberine complex extended over a duration of 50 ns. To analyze the stability and flexibility of the system, Root Mean Square Deviation (RMSD) analysis, along with the assessment of the Root Mean Square Fluctuation (RMSF) of residues, were performed using the Xmgrace software package. These analyses provided insights into the conformational changes and dynamic behavior of the complex. Furthermore, the binding free energy between the TatB and berberine was quantitatively determined through the application of the MM-PBSA (Molecular Mechanics Poisson-Boltzmann Surface Area) method. This approach enabled the estimation of the thermodynamic stability of the complex. Ultimately, the most stable configuration of TatB-berberine was analyzed using Discovery studio, contributing significantly to the understanding of the underlying interaction mechanisms.

### 2.15 Statistics

Two-tailed, paired, student’s t test p-value (GraphPad Prism) calculations determined statistical significance as detained in each figure legend and **Supplemental Table 1**.

## 3. RESULTS

### Descriptions of the compounds used in this study

All the three compounds – Berberine (BBR), isoliquiritigenin (ILG), and 4′-dihydroxyflavone (DHF) are in powder forms. Characteristically, BBR is yellow in color. HPLC analysis of BBR fingerprint showed the peak at retention time 6.2 min with ***λ*** max at 264 nm (**Figure 1A**). DHF is light yellow in color and HPLC fingerprint showed the peak of DHF at retention time of 5.8min with ***λ*** max at 330 nm (**Figure 1B**). ILG is also yellow in color and HPLC fingerprint showed the peak of ILG is at the retention time of 7.3min with ***λ*** max at 364 nm (**Figure 1C**). The purity of each compound was >95%. The heavy metal levels in these compounds are less than 20 ppm. Thus, the purity and quality of BBR, DHF, and ILG comply with the United States Pharmacopeia (USP) standard.

**Figure 1:**
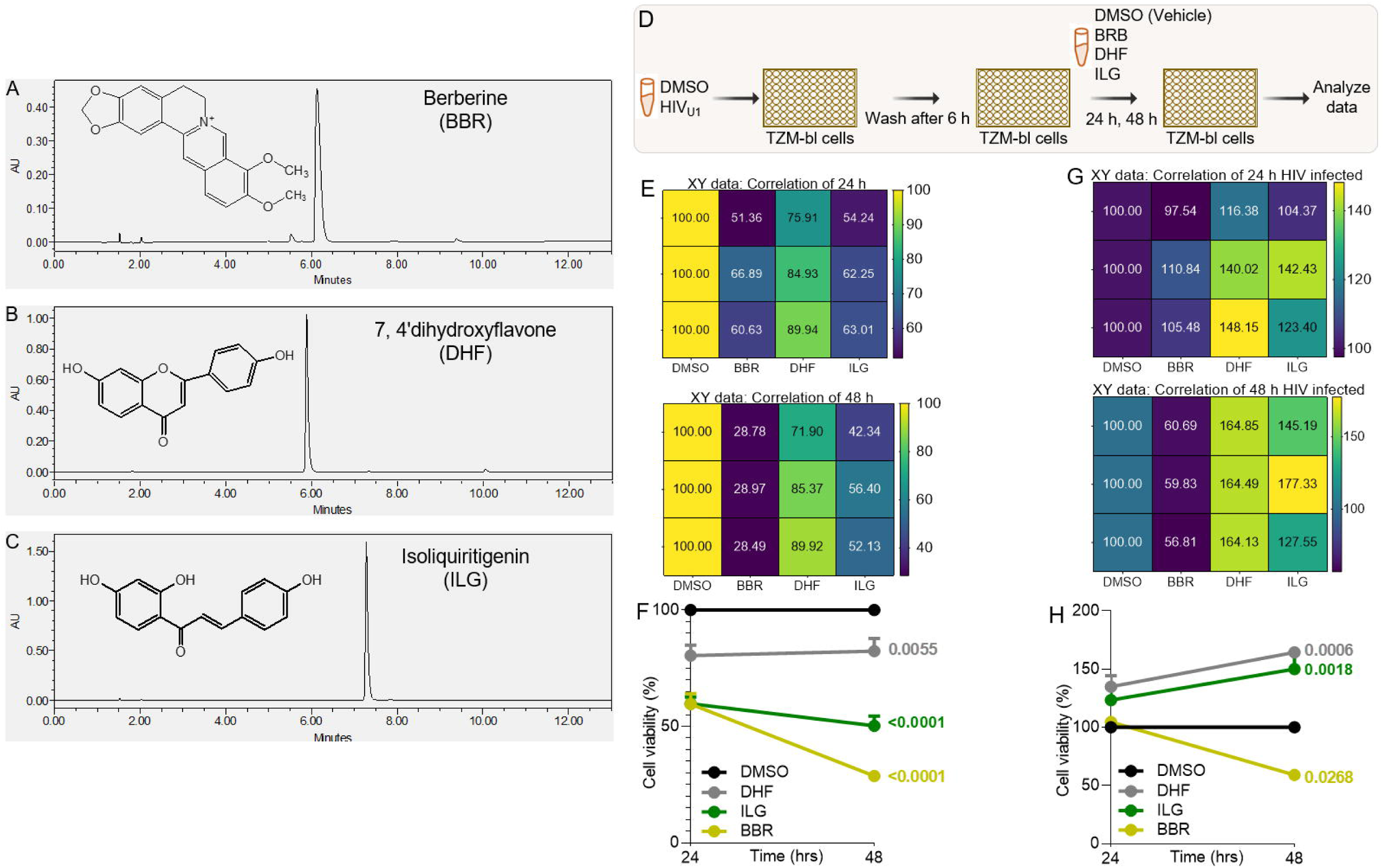
Description of TCM compounds and their effects on the viability of TZM-bl cervical cancer cell line infected or not infected with HIV: **A - C)** Chemical structures and HPLC fingerprints of **A)** BBR, **B)** DHF, **C)** ILG. **D)** Infographic of experimental protocol where cells were infected and treated 6 hours after infection with readout at 24 hours and 48 hours post infection. **E, F)** Results of the viability of uninfected TZM-bl cells treated with DMSO or the different TCMs over time (24 h, 48 h) displayed as **E)** correlation matrix showing the direction and strength of the TCMs on cell viability, **F)** % viable cells. **G, H)** Results of the viability of HIV infected TZM-bl cells treated with DMSO or the different TCMs over time (24 h, 48 h) displayed as **G)** Correlation matrix showing the direction and strength of the TCMs on cell viability, **H)** % viable cells. Experiments were repeated at least three times. Statistical differences were assessed by GraphPad Prism Mixed Effects Models using Two-stage linear step-up procedure of Benjamini, Krieger and Yekutieli.

### The effects of TCM compounds on the viability of TZM-bl cervical cancer cells is dependent on HIV infection status of the cell

Cervical cancer cell viability inhibition potential of three TCM – berberine (BBR),4′-dihydroxyflavone (DHF), and isoliquiritigenin (ILG) was measured in TZM-bl cells as indicator cell line using the MTT method. Cell viability was assessed after the cells were infected with HIV_U1_ [40] or left uninfected and treated with 50 µl of 0.1X DMSO used as vehicle (control) or 40 µg of compounds in 50 µl of media (**Figure 1D**). In uninfected TZM-bl cells, viability was inhibited by all three TCM compounds (**Figure 1E left, 1F**). Prolongation of duration of treatment resulted in increased inhibition by ILG and BBR, with BBR being the most potent inhibitor at 48 hours post treatment (**Figure 1E right, 1F**). Subsequently, the effect of the compounds on the viability of HIV infected cells when added early in infection was measured (**Figure 1D**). In HIV infected cells, ILG and DHF enhanced cell viability both at 24 hours and 48 hours while BBR has no effect of cell viability 24 hours after treatment (**Figures 1G, 1H**). However, by 48 hours BBR significantly and potently inhibited cell viability (p=0.0268) in contrast to ILG (p=0.0006) and DHF (p=0.0018) that significantly increased cell viability (**Figure 1G right, 1H**).

### TCM compounds have distinct effects on HIV infected cervical cancer cell viability

To examine the effect of the three TCM compounds on HIV infection of cervical cancer cells that mirrors more of natural infection where infection was allowed to proceed for 24 hours. Cells were left uninfected or infected with HIV and then treated with DMSO or BBR, DHF, ILG (**Figure 2A**). The effects of the BBR, DHF, ILG on HIV promoter transactivation were determined 24 hours later by measuring promoter activity using the Steady-Glo^®^ (Promega) luminescence assay as previously described [39]. BBR, DHF, ILG inhibited HIV LTR promoter transactivation by ∼67 %, ∼48 %, ∼66 %, respectively in the luciferase assay (**Figure 2B, C**). Since luciferase assay only assesses HIV promoter activity and not HIV RNA, we used RT-qPCR to assess the effect of the TCM on cell-associated HIV RNA (CA-HIV RNA). Similar to its effect on HIV LTR promoter activity, BBR, DHF, ILG inhibited CA-HIV RNA by ∼68 %, ∼20 %, ∼35 %, respectively (**Figure 2D, E**). Additionally, to quantify the amount of HIV released into the supernatants following treatment with the different TCMs, we used the HIV reverse transcriptase (RT) assay [37, 38, 45] to analyze extracellular RT (exRT) as a proxy to determine the concentration of HIV in the various supernatants. As shown in the correlation matrix (**Figure 2F**), BBR and ILG decreased the amounts of exRT compared to DMSO while DHF did not alter HIV RT levels. However, while BBR and ILG significantly decreased the amounts of exRT, BBR had a greater and significant effect in decreasing exRT compared to ILG (**Figure 2G).** Since BBR showed more potent inhibition of HIV promoter activity (**Figure 2B, C**), CA-HIV RNA expression (**Figure 2D, E**), and exRT levels (**Figure 2F, G**) we focused on BBR in subsequent experiments.

**Figure 2:**
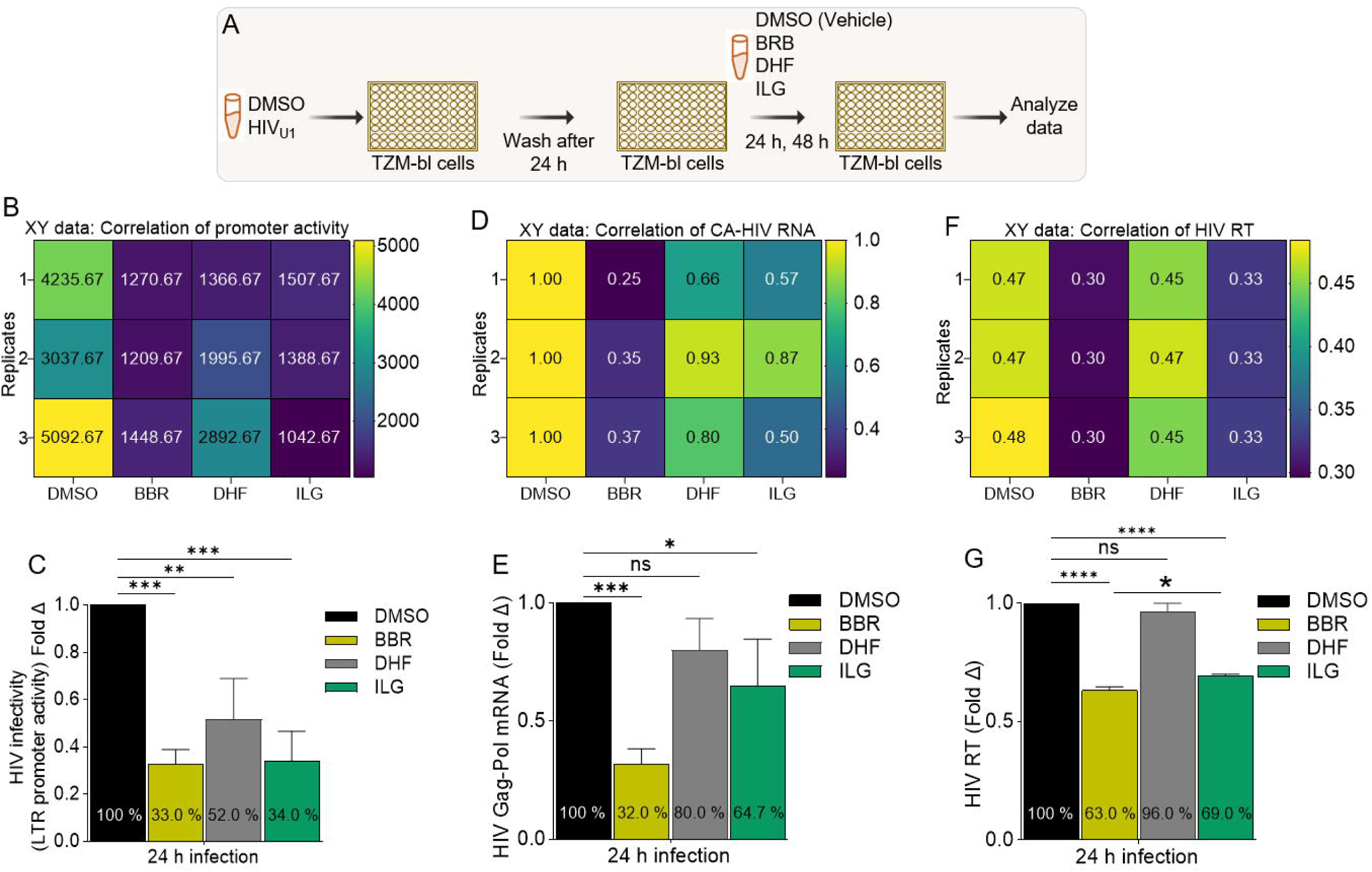
TCM compounds have distinct effects on HIV infected cervical cancer cell viability: **A**) Infographic of experimental protocol where cells were infected, and infection allowed to proceed for 24 hours prior to treatment and readout 24 hours later. **B, C**) Results of the HIV LTR promoter transactivation in TZM-bl cells treated with DMSO or the different TCMs for 24 h displayed as **B**) Correlation matrix showing the direction and strength of the TCMs on HIV LTR promoter activity, **C**) Fold change. **D, E**) Results of cell-associated HIV RNA (CA-HIV RNA) in TZM-bl cells treated with DMSO or the different TCMs for 24 h displayed as **D**) Correlation matrix showing the direction and strength of the TCMs on HIV LTR promoter activity, **E**) Fold change. **F, G**) Results of the production of extracellular HIV reverse transcriptase (exRT) released by infected TZM-bl cells treated with DMSO or the different TCMs for 24 h displayed as **F**) correlation matrix showing the direction and strength of the TCMs on HIV LTR promoter activity, **G**) Fold change. Experiments were repeated at least three times. Statistical differences were assessed by GraphPad ordinary one-way ANOVA with Šídák’s multiple comparisons test. **** p < 0.0001, *** p < 0.0002 – 0.0005, ** p < 0.022, * p < 0.02, and ns = non-significant.

### Cancer cells internalize BBR and exhibit morphological change

To further examine the effect of TCM BBR on HIV infection of cervical cancer cells, TZM-bl cells were left uninfected or infected with HIV and then treated with DMSO or 40 µg BBR compounds (**Figure 3A**). Since BBR naturally fluoresces, we used the fluorescent properties to show that by 1 hour post treatment, BBR is internalized by TZM-bl cells (**Figure 3B**) and quantitatively significant (**Figure 3C**). Because clustering of cancer cells is a pathophysiological process that depicts increased malignancy and HIV infection may promote cell clustering, TZM-bl cells were also assessed for morphological changes, including cell to cell clustering. Compared to uninfected cells, HIV infection resulted in higher levels of cell clustering (**Figure 3D, compare top and middle**). Infected cells treated with BBR exhibit obvious morphological changes, such as smaller size, rounded shape, and decreased levels of cell clustering compared to HIV infected cells treated with DMSO (**Figure 3D, compare middle and bottom**). Together, these findings suggest that BBR may diminish HIV-induced cell clustering in cervical cancer cells, as a potential mechanism of inhibiting molecular signaling and cell to cell virus spread.

**Figure 3:**
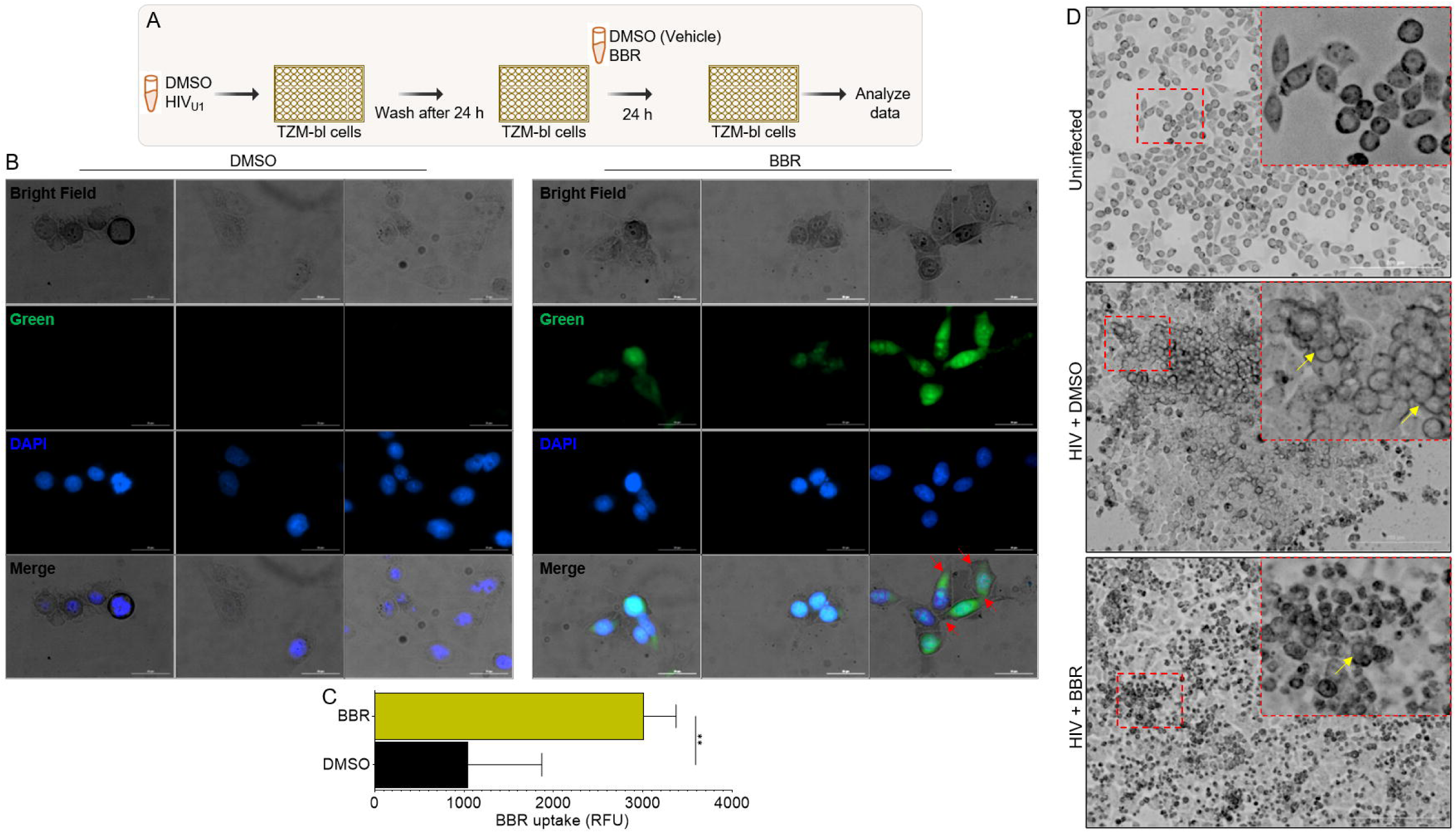
Cancer cells internalize BBR and BBR induces morphological changes in the cells: **A**) Infographic of experimental protocol where cells were infected, and infection allowed to proceed for 24 hours prior to treatment and readout 24 hours later. **B**) Representative images of HIV infected TZM-bl cells treated with BBR or DMSO for 24 hours and stained with DAPI. Images were captured with LionHeart 60X objective and analyzed using the Gen5 software. Green = BBR, blue = DAPI. Red arrows indicate areas of cell-to-cell contacts. **C**) BBR uptake was analyzed by quantifying 3 fields of view with Gen5 software and presented as raw relative florescence units (RFU). **D)** Morphology of uninfected TZM-bl cells (top), HIV-induced TZM-bl clustering (middle), and BBR-mediated inhibition of HIV-induced TZM-bl clustering (bottom). Red squares are the enlarged areas showing clustered cells and presented as inset within each image. Experiments were repeated at least three times. Statistical differences for pane C was assessed by GraphPad’s unpaired t test with Welch’s correction. ** p < 0.0012.

### BBR induce correlated inhibition of multiple processes in the HIV infection cycle

To further assess the potency of BBR on HIV infection, we conducted a concentration dependent study of BBR in TZM-bl cells that were infected with HIV then treated with DMSO or BBR (**Figure 4A**). BBR elicited a concentration (0, 20, 40, 80, 100 µg) dependent effects in cell viability, HIV LTR promoter activation, and CA-HIV RNA expression (**Figure 4B**, Tukey’s multiple comparisons test presented in Table below). Correlation matrix analysis was used to show the direction of BBR concentrations on cell viability (**Figure 4C**). We found that 20 µg of BBR led to ∼112 % increase in cell viability. In contrast, increased concentration of BBR 40, 80, 100 µg led to ∼86, ∼83, ∼51 % decrease in cell viability (**Figure 4B, 4C**). In parallel, BBR showed concentration dependent decrease in HIV LTR promoter activity as shown by the graph and correlation matrix (**Figure 4B, 4D**). While 20 µg of BBR did not significantly (∼81 %) decrease HIV LTR promoter activity, 40, 80, 100 µg of BBR respectively led to ∼53, ∼44, ∼34 % decrease in HIV LTR promoter activity (**Figure 4B, 4D**). Additional analysis of CA-HIV RNA expression also revealed a significant decrease in CA-HIV RNA (**Figure 4E**), where 20, 40, 80, 100 µg of BBR led to ∼61, ∼83, ∼96, ∼98 % decrease in CA-HIV RNA expression (**Figure 4B, 4E**). To provide context for the ability of BBR to inhibit cell viability and HIV infection, we evaluated the EC_50_ (https://www.aatbio.com/tools/ec50-calculator) of BBR [46]. We found that BBR exhibit dose-dependent correlated inhibition of cell viability, HIV LTR promoter activity, and CA-HIV RNA expression with EC_50_ of 90.3788 µg, 29.0839 µg, and 23.0632 µg respectively (**Figure 4F**).

**Figure 4:**
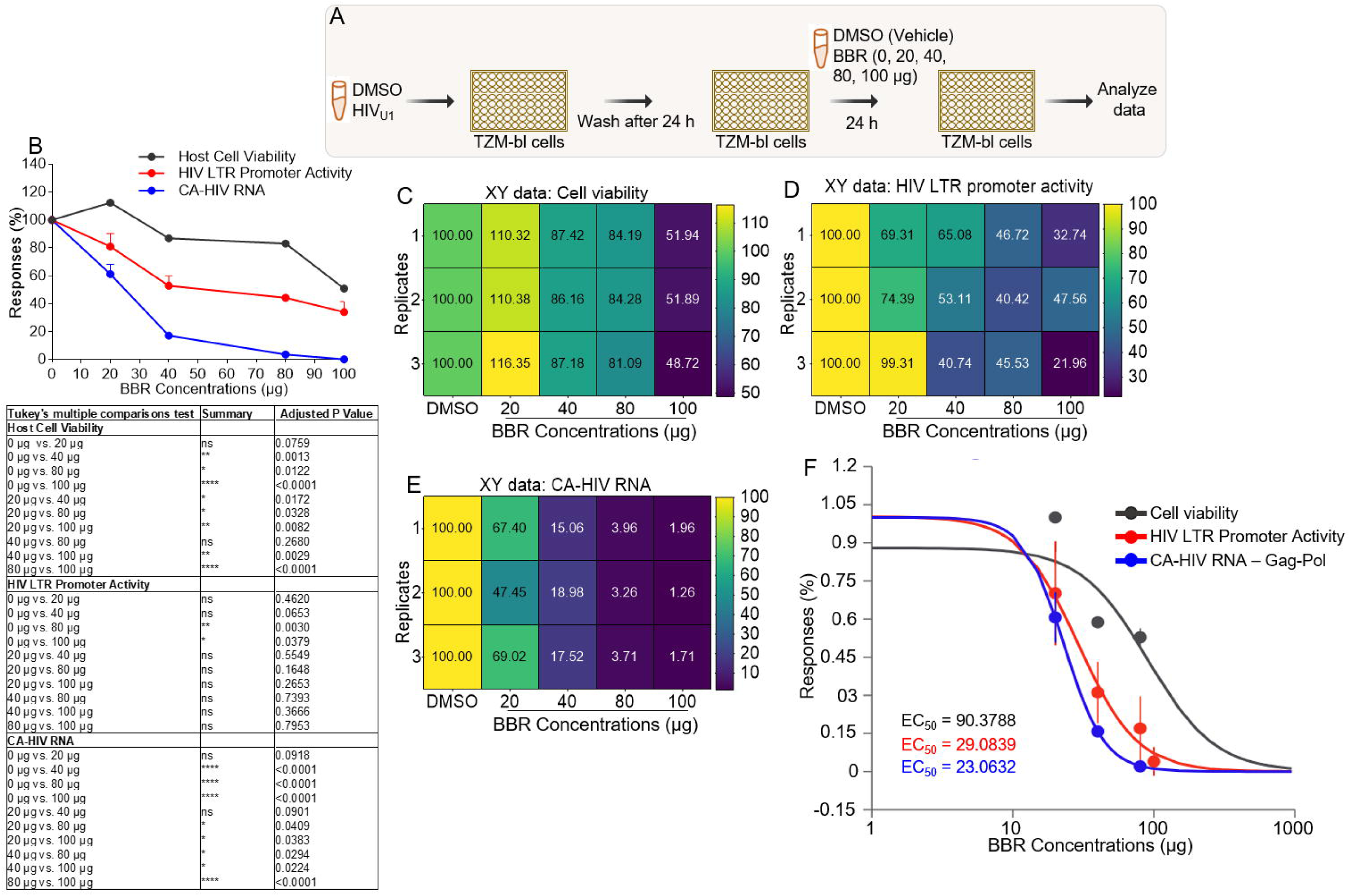
BBR induces correlated inhibition of multiple processes in the HIV infection cycle: **A**) Infographic of experimental protocol where cells were infected, and infection allowed to proceed for 24 hours prior to treatment and readout 24 hours later. **B**) XY plot of BBR concentration (0, 20, 40, 80, 100 µg) versus host responses (t**op**) with table of GraphPad Prism’s statistical analysis (Mixed effects analysis, Tukey’s multiple comparisons test, **bottom**). **C-E**) Correlation matrix showing the direction and strength of BBR on **C**) cell viability., **D**) HIV LTR promoter activity, **E**) CA-HIV Gag-Pol RNA. **F**) The EC_50_ of BBR determined using an online calculator (https://www.aatbio.com/tools/ec50-calculator). Experiments were repeated at least three times. Statistical differences were assessed by GraphPad ordinary one-way ANOVA with Šídák’s multiple comparisons test and unpaired t test with Welch’s correction. **** p < 0.0001, *** p < 0.0002 – 0.0005, ** p < 0.022, * p < 0.02, and ns = non-significant.

### BBR inhibits correlated HIV infection and migration of HIV infected cervical cancer cells

We have thus far shown that BBR synchronizes inhibition of cell viability and HIV infection. Since cell migration is a hallmark of cancer that promotes metastasis and migration of HIV infected cells from the genital mucosa to lymphoid organs is key to systemic viral spread, we assessed the effect of BBR on migration of HIV infected cells. Using our previously described wound healing assay [43, 47], we examined the migration of uninfected and HIV infected TZM-bl cells in response to the mechanical scratch wound in the absence or presence of different concentrations of BBR (**Figure 5A**). Representative images of the scratch areas from time points 0, 18, and 24 hours are shown in **Figure 5B**. Compared to uninfected and untreated cells with ∼69% and ∼59% wound area respectively at 18 and 24 hours, HIV infected and untreated cells had ∼55% and 49% wound area at the same time points (**Figure 5C, left**). These data indicate that HIV promotes migration (wound closure) of cervical cancer cells. We further show that in the presence of 40 µg of BBR, HIV infected cells showed time-dependent increase in cell migration with ∼38% wound area compared to ∼59% migration in uninfected cells treated with the same concentration of BBR at 24 hours (**Figure 5C, center**). Interestingly, 100 µg of BBR inhibits migration of both uninfected and HIV infected cells with wound areas of ∼68% and 70% respectively, with no infection-dependent differences (**Figure 5C, right**). To link inhibition of HIV infection and migration of infected cells by BBR, we analyzed the levels of HIV exRT in supernatants collected from the wells of the migration assay. As expected, BBR suppressed the levels of HIV exRT from 100 % to ∼66 % and 64 % for 0 µg, 40 µg, and 100 µg of BBR respectively (**Figures 5D, E**). These data suggest that BBR has a concentration-dependent effect on HIV infection and migration of HIV infected cervical cancer cells with 100 µg BBR concentration delivering the most effect.

**Figure 5:**
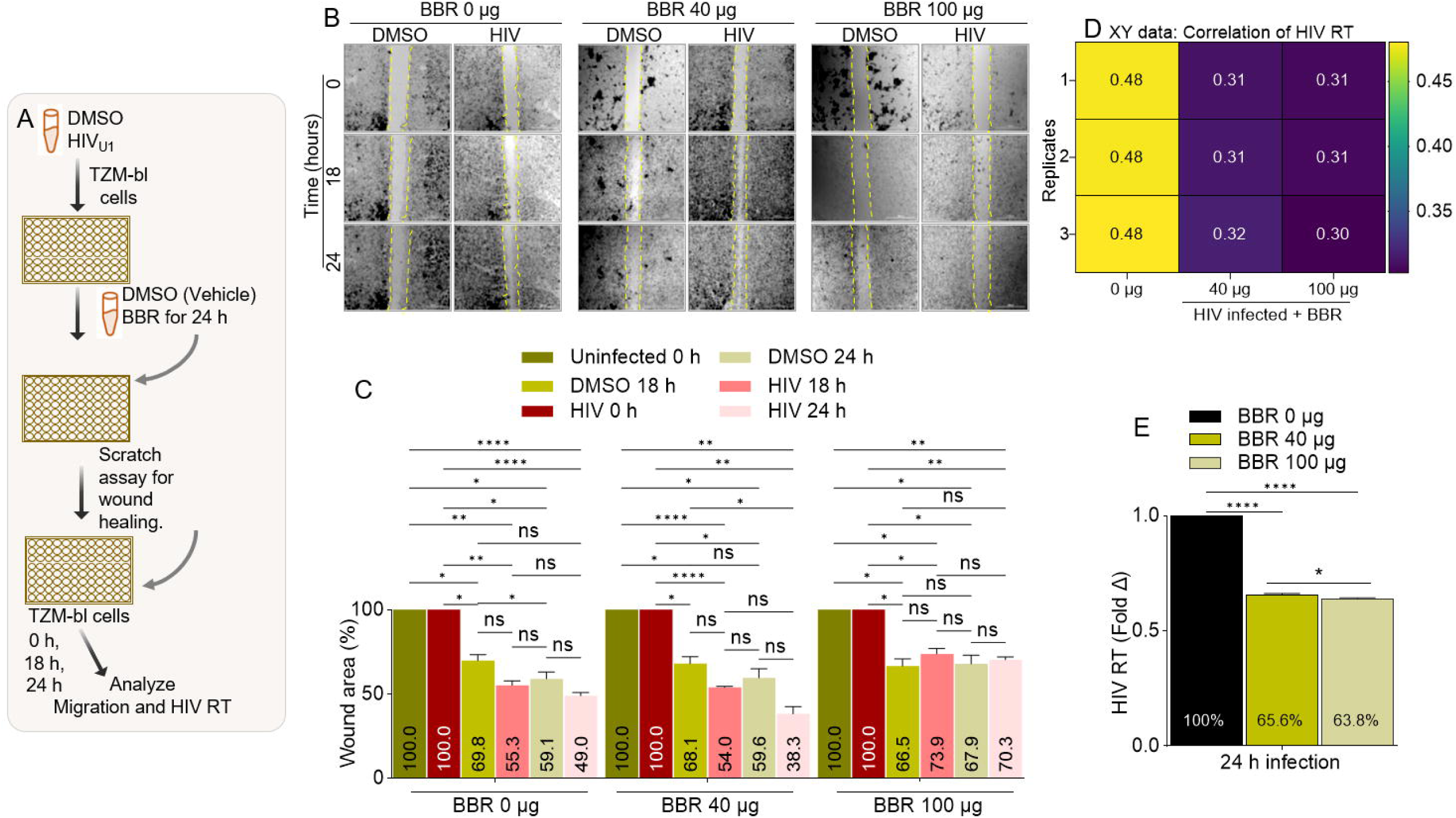
BBR inhibits correlated HIV infection and migration of HIV infected cervical cancer cells: **A**) Infographic of experimental protocol. **B**) Representative images of wound tracks and wound areas. The vertical broken lines track the wound areas. Images were captured with LionHeart 4X objective in a 6×6 montage to display all frames of a multi-frame image array to enable visualization of the areas. Scale = 3000 µm. **C**) Time and concentration dependent plot of wound area. **D, E**) Levels of HIV exRT secreted into the supernatants of cells in migration (wound healing) chamber presented as **D**) Correlation matrix showing the direction and strength of BBR on HIV exRT. **E**) Fold change in HIV exRT levels. Experiments were repeated at least three times. Statistical differences were assessed by GraphPad’s 2way ANOVA with Tukey’s multiple comparisons test for panel C, and ordinary one-way ANOVA with Šídák’s multiple comparisons test and unpaired t test with Welch’s correction for panel E. **** p < 0.0001, ** p < 0.0032, * p < 0.018 – 0.03, ns = non-significant.

### BBR inhibits Tat-induced HIV LTR promoter transactivation and Tat mRNA expression

Transcription of HIV is governed by the viral LTR, and the primary function of Tat is to transactivate HIV LTR promoter [48]. Thus, if BBR affects HIV Tat function, it should strongly inhibit HIV LTR-directed promoter transactivation induced by Tat. We used our previously described protocol to investigate the effect of BBR on Tat-induced HIV LTR promoter transactivation [39]. Thus, TZM-bl cells were either infected with HIV_U1_ or treated with recombinant Tat (rTat) for 24 hours followed by treatment with DMSO or BBR (**Figure 6A**). While BBR alone did not significantly alter HIV LTR promoter activation as show by the correlation matrix (**Figure 6B**) and bar graph (**Figure 6C**), cells infected with HIV or treated with rTat exhibited significant increase in the levels of luciferase activity, a marker of HIV LTR promoter activation, compared to untreated cells (**Figure 6D, 6E)**. BBR significantly suppressed HIV (**Figure 6F, 6G**) and rTat (**Figure 6H, 6I**). Since Tat directly controls HIV replication, we examined the effect of BBR on Tat mRNA expression in infected cells. The level of CA-HIV Tat mRNA is dependent on BBR concentration, where 40 µg increased CA-HIV Tat RNA by ∼347 % while 100 µg suppressed CA-HIV Tat RNA to ∼46% (**Figure 6J, left**). Furthermore, there is a dose-dependent decrease in the mRNA of Gag-Pol from ∼22% to 6% with 40 µg and 100 µg respectively (**Figure 6J, bottom**), suggesting that BBR may have a concentration dependent effect on the expression of CA-HIV RNA species. As expected, BBR decreased cell viability of uninfected and infected cells in a dose dependent manner (**Figure 6K, L**). Although, the effect of BBR is more robust in uninfected cells compared to their infected counterparts (**Figure 6K, L**).

**Figure 6:**
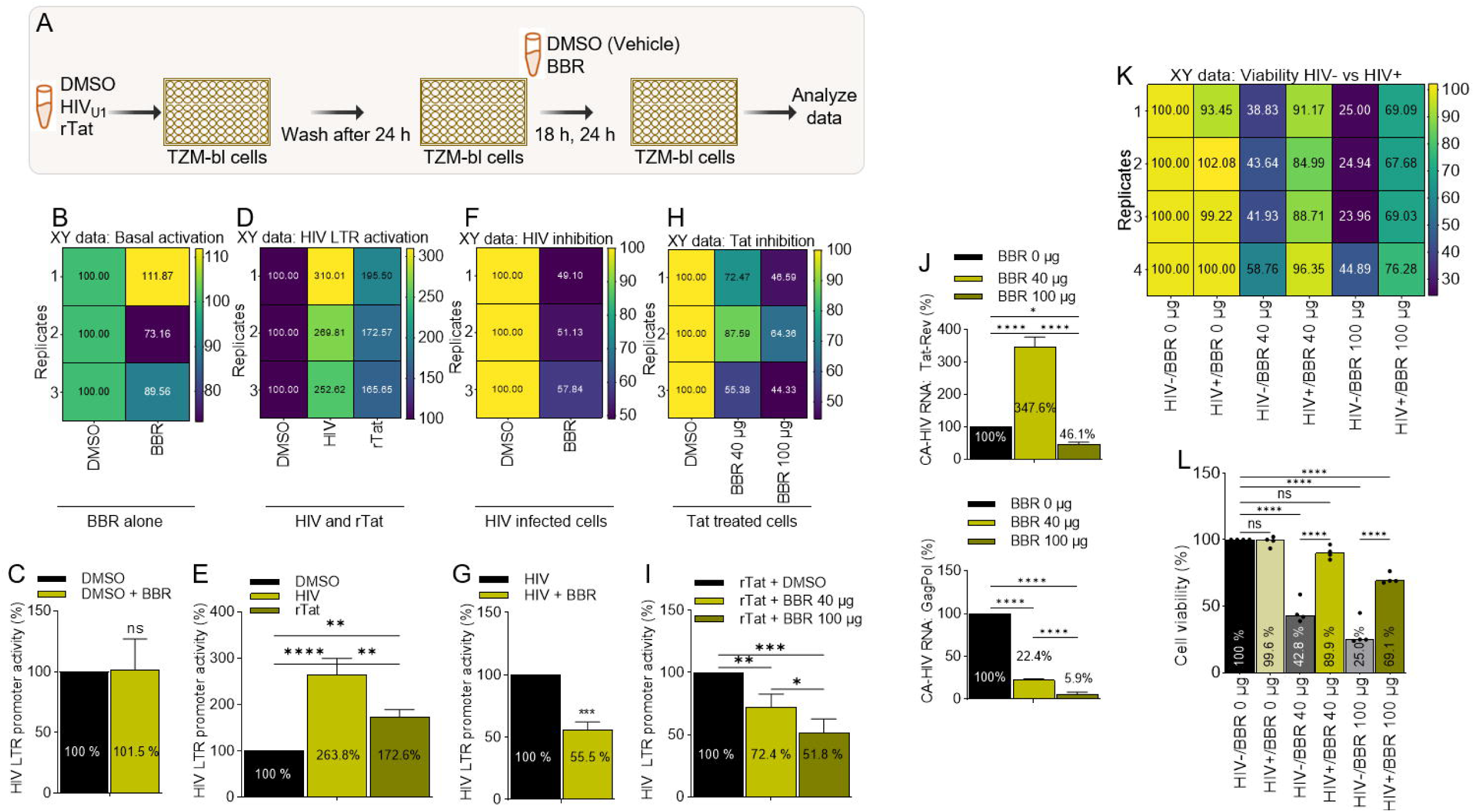
BBR inhibits Tat-induced HIV LTR promoter transactivation and Tat mRNA expression: **A**) Infographic of experimental protocol. **B, C**) HIV LTR promoter activity in uninfected TZM-bl cells treated with DMSO or 40 µg of BBR **B**) correlation matrix showing the direction and strength of the BBR on HIV LTR promoter activity, **C**) Percent (%) luminescence as readout of promoter transactivation. **D, E**) HIV LTR promoter activity in uninfected DMSO treated, HIV infected, or rTat treated cells TZM-bl, **D**) Correlation matrix showing the direction and strength of the BBR on HIV LTR promoter activity, **E**) Percent (%) luminescence as readout of promoter transactivation. **F, G**) HIV LTR promoter activity in HIV infected DMSO or BBR treated, cells, **F**) Correlation matrix showing the direction and strength of the BBR on HIV LTR promoter activity, **G**) Percent (%) luminescence as readout of promoter transactivation. **H, I**) HIV LTR promoter activity in TZM-bl cells treated with DMSO, rTat, or different concentrations (40, 100 µg) of BBR. **H**) Correlation matrix showing the direction and strength of BBR on HIV LTR promoter activity, **I**) Percent (%) luminescence as readout of promoter transactivation. **J**) CA-HIV MS Tat RNA, (top) and US Gag-Pol RNA (bottom). **K**) Correlation matrix showing the direction and strength of different concentrations (0, 40, 100 µ) of BBR on uninfected (HIV-) and HIV infected (HIV+) TZM-bl cell viability. **L**) Percent (%) cell viability. Experiments were repeated at least three times. Statistical differences were assessed by GraphPad ordinary one-way ANOVA with Šídák’s multiple comparisons test and unpaired t test with Welch’s correction. **** p < 0.0001, *** p < 0.0002 – 0.0005, ** p < 0.022, * p < 0.02, and ns = non-significant.

### BBR inhibits Tat-induced migration and matrix invasion of cervical cancer cells

The oncogenic activities of HIV is driven in part by the Tat protein, which may depend on the ability of Tat to promote cell motility [21, 23]. As a result, we assessed the effect of BBR on Tat-induced migration and matrix invasion of cervical cancer cells (**Figure 7A**). Tat expression plasmid transfected into TZM-bl cells transactivated HIV LTR promoter while BBR at different concentrations (40 µg or 100 µg) did not transactivated HIV LTR promoter (**Figure 7B**). However, BBR significantly suppressed Tat-mediated HIV LTR promoter transactivation (**Figure 7C**). The effect of Tat on cell migration was assessed using the scratch (wound healing) assay. Representative images of the scratch areas from time points 0, 18, and 24 h are illustrated in **Figure 7D**. TZM-bl cervical cancer cells efficiently migrated to close the wound area with 64.6 % and 50.5 % wound area remaining at 18 and 24 hours respectively. In contrast, the presence of pTat promotes cell migration with 44.7 % and 21.5 % wound area remaining at 18 and 24 hours respectively (**Figure 7E**). As expected, BBR inhibits basal TZM-bl cell migration in a concentration dependent manner with 78.1 % and 70.7 % wound area remaining at 18 and 24 hours respectively for BBR 40 µg; and 91.7 % and 88.3 % for BBR 100 µg respectively (**Figure 7E**). Interestingly, BBR significantly (**Supplemental Table 1**) inhibits Tat-mediated TZM-bl cell migration in a concentration dependent manner with 44.7 %, 76.6 %, 98.0 % wound area remaining at 18 hours in the presence of 40 µg of BBR and 21.5 %, 56.9 %, 93.4 % wound area remaining at 24 hours in the presence of 100 µg of BBR (**Figure 7E**). Similar to migration, TZM-bl cancer cells significantly invade matrix by breaking through Matrigel barrier in a transwell invasion assay (**Figure 7A**). The representative images of the membrane insert stained with Giemsa (**Figure 7F**) and the number of quantified membrane-bound cells (**Figure 7G**) show that compared to DMSO treated cells, (100 %) pTat increased the number of membrane-bound cells (237%) and BBR at 40 µg and 100 µg significantly decreased the number of membrane-bound cells and inhibited Tat-mediated invasion in a concentration dependent manner (**Figures 7F, 7G**). Further analysis of the effect of BBR on Tat-mediated cell invasion by assessing the number of cells that completely invaded and settled in the basal chamber support the observation that BBR potently inhibit TZM-bl invasion and block Tat-mediated cell invasion in a concentration dependent manner (**Figures 7H, 7I**).

**Figure 7:**
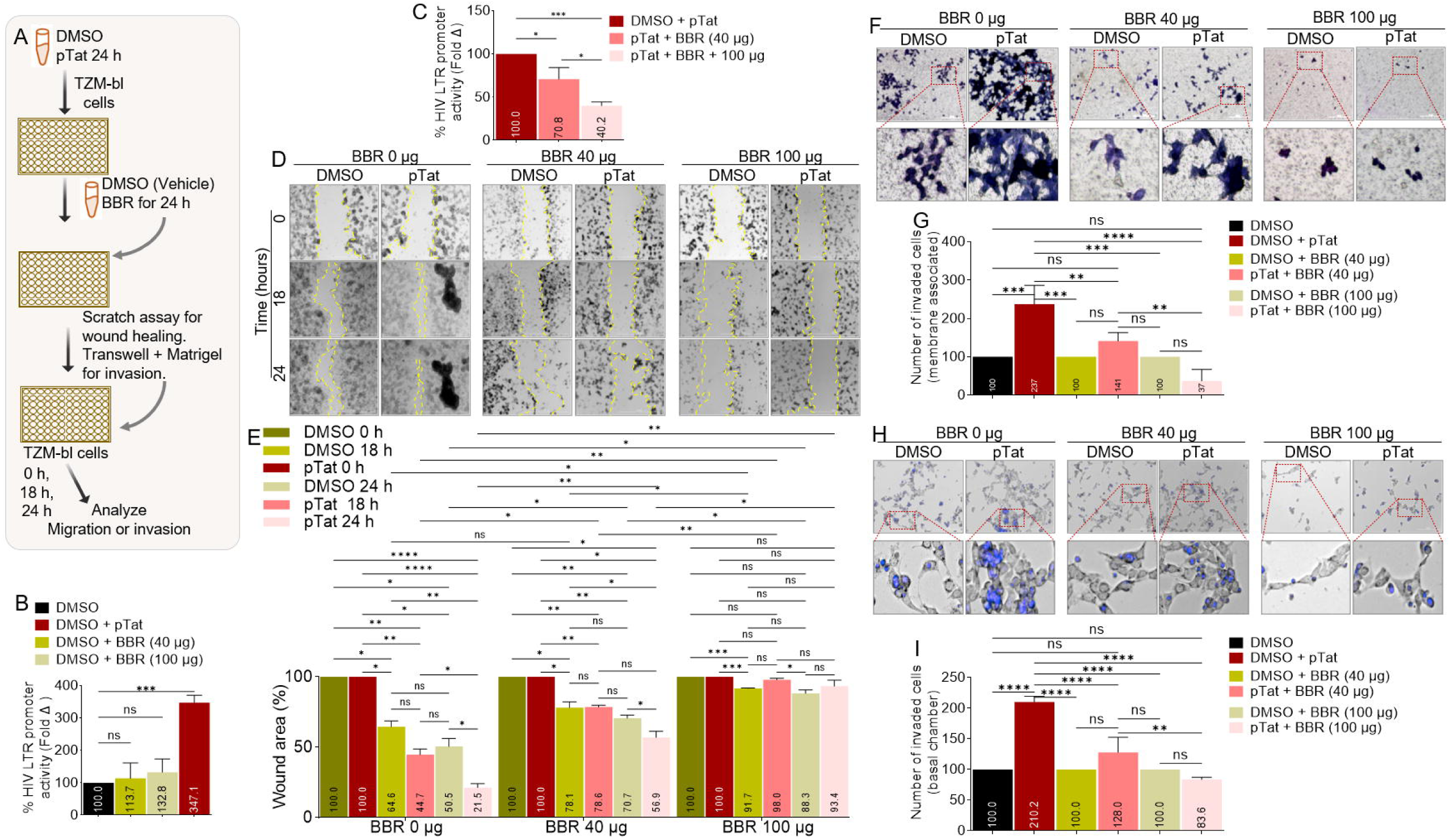
BBR inhibits HIV Tat-mediated migration and matrix invasion of cervical cancer cells: **A**) Infographic of experimental protocol. **B**) The effects of BBR and pTat on HIV LTR promoter transactivation 24 hours post transfection. **C**) The effects of BBR on pTat-mediated HIV LTR promoter transactivation 24 hours post transfection. **D**) Representative images of wound tracks and wound areas. The vertical broken lines track the wound areas. Images were captured with LionHeart 4X objective in a 6×6 montage to display all frames of a multi-frame image array to enable visualization of the areas. Scale = 3000 µm. **E**) Time and concentration dependent plot of wound area. **F**) Representative images of cells that invaded Matrigel-coated membrane insert stained with Giemsa. **G**) The number of quantified membrane-bound cells is shown in panel **F**. **H**) Representative images of cells that invaded through Matrigel into the basal chamber of the transwell and stained with NucBlue. **I**) The number of quantified basal chamber cells in panel **H**. The red squares in panels **F** and **H** represent the enlarged areas below each panel. Experiments were repeated at least three times. Statistical differences were assessed by GraphPad’s 2way ANOVA with Tukey’s multiple comparisons test for panel E, ordinary one-way ANOVA with Šídák’s multiple comparisons panels B, C, G, I. **** p < 0.0001, ** p < 0.0032, * p < 0.018 – 0.03, ns = non-significant. Complete statistical data for panel **E** is presented in **Supplemental Table 1.**

### BBR binds HIV Tat amino acid residues through non-covalent interaction

Since BBR inhibits the activities of HIV and HIV Tat, we sought to elucidate the molecular mechanisms using molecular docking and dynamic stimulation. It is widely acceptable that the strength of binding is defined by negative energy value. Our docking analysis initially identified a binding energy of - 6.4 kcal/mol and −5.9 kcal/mol at 0 ns for TatB-BBR and TatC-BBR interactions (**Table 2**), indicating a binding possibility between the two molecules. However, following a 50 ns molecular dynamics simulation, the average binding free energy, as calculated using the MM-PBSA method, was −11.54 kcal/mol and −8.6 kcal/mol for TatB-BBR and TatC-BBR respectively, indicating a strong and stable binding between TatB and BBR. This marked reduction (**Table 3**) underscores the enhanced stability of TatB-BBR complex, but not so much of TatC-BBR upon undergoing molecular dynamics simulation. The most favorable and stable configuration attained by the TatB-BBR complex is depicted in 2D (**Figure 8A**) and 3D (**Figure 8B**) formats. The dynamic stimulation analysis revealed that BBR establishes π-amide stacking with Tat LYS71, which significantly contributes to stabilizing the TatB-BBR complex. Additionally, alkyl interactions were observed between BBR and Tat amino acid residues CYS22, LYS28, VAL67, and PRO73, further reinforcing the TatB-BBR complex stability. Moreover, the cysteine residues CYS27 and CYS34 support the TatB-BBR complex through π-sulfur interactions. Lastly, carbon-hydrogen bonds were formed between BBR and the amino acids GLN17, LEU69, LYS28, and GLN71, providing additional stability to the TatB-BBR complex. The RMSD (Root Mean Square Deviation) values for TatB and BBR are presented in **Figure 8C**. Initially, the RMSD of TatB increases, and subsequently reaches a stable equilibrium during 30-50 ns, whereas the RMSD of BBR remains relatively constant throughout the entire 50 ns period. Additionally, the Root Mean Square Fluctuation (RMSF) analysis was performed to assess the flexibility of TatB residues, as depicted in **Figure 8D**. Importantly, the active site residues displayed low flexibility values, indicating that the active site remains stable and does not undergo substantial conformational changes during the simulation. This suggests that the binding of BBR to TatB does not disrupt the structural integrity of the active site of TatB.

**Figure 8:**
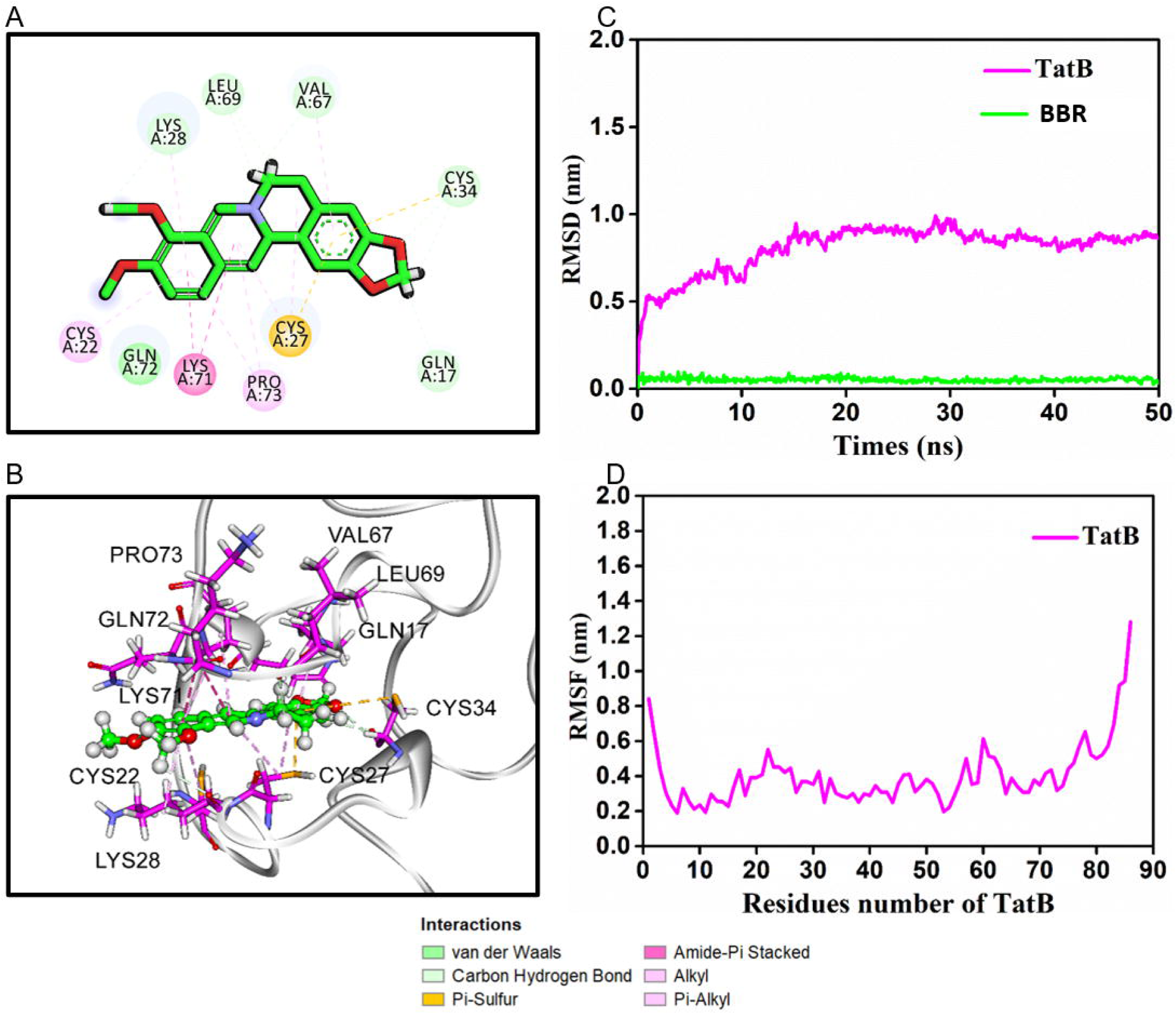
BBR binds HIV Tat amino acid residues through non-covalent interaction: A, **B**) Binding explorations of TatB-BBR complex. Molecular dynamic simulations predict lowest-energy binding mode of TatB-BBR for **A**) 2-dimentional (2D) and **B**) 3-dimentional (3D). **C**) RMSD values for TatB and BBR over 50 ns. **D**) RMSF of the TatB protein residues. For resident proteins, carbon, oxygen and nitrogen are highlighted in purple, red and blue, respectively. For berberine, the green, red, and blue stand for C, O and N, respectively.

## 4. DISCUSSION

A significant finding of this study is the remarkable ability of BBR to potently and correlatively decrease viability of TZM-bl cervical cancer cells, inhibit expression of CA-HIV RNA (Tat-Rev, Gag-Pol) species, block Tat mediated HIV LTR promoter transactivation, and inhibit HIV Tat-mediated migration and matrix invasion of HIV infected TZM-bl cells.

BBR is an isoquinoline alkaloid small molecule compound present in the roots, rhizomes, stem barks, and leaves of various herbs and plants of the Berberis genus [49], and can be obtained from plants or chemically synthesized. Characteristically, BBR powder is yellow, odorless, and alkaloid bitter [50]. BBR (C_20_H_18_NO_4_) is a quaternary ammonium salt of an isoquinoline alkaloid (PubChem CID: 2353) with a molecular weight of 336.4 g/mol [49]. Its solubility is poor, as it is minimally soluble in water, and slightly soluble in ethanol or methanol [50]. A review of multiple studies evaluating various TCMs, including BBR for the treatment of viral infections and their associated cancers, such as cervical cancer [51], highlight increasing focus on their potential use in the treatment of viral infections and their associated cancers. In addition to its anti-viral qualities, BBR has lipid-lowering, anti-diabetic, anti-inflammatory properties [32–34, 52–55].

In this study, we showed that BBR is a promising natural product with inhibitory effects on the viability of cervical cancer cells infected with HIV or treated with HIV Tat protein. Our findings that BBR inhibits HIV infection in cervical cancer cells agree with studies demonstrating the antiviral effects of BBR. For example, BBR has been shown to inhibit HIV RT activity [56], protect cells against Epstein–Barr virus and its associated tumors [57, 58], target Kaposi’s sarcoma-associated herpesvirus and its associated cancers [59], as well as inhibits HSV 1, HSV 2 [60–62], and HCMV [63] infections. Our data support the reported effect of BBR on HIV RT because we found that cells treated with BBR secrete significantly reduced exRT (**Figures 2F, G**; **5D, E**).

Although BBR is sparingly soluble in water, we showed that TZM-bl cells internalized BBR, supporting a previous report by Pang et al., [64]. Uptake of BBR by cells may help improve its efficacy and overcome the insolubility shortfall. In TZM-bl cells, HIV infection increased cell clustering, but BBR altered the cell to cell contact behaviors between infected cells. Since cell clustering involves membrane rearrangements that may promote HIV cell to cell virus transmission and cancer progression, and may link viruses to cancer, we speculate that BBR may prevent cancer induced cell to cell signaling and chromosomal instability that may underlie malignant properties of cervical cancer. Of note, it has been suggested that cell clustering may enhance energy metabolism of mesenchymal tumor cells to sustain their proliferation and invasion [65].

Our data showed that BBR exhibits dose-dependent effect on cell viability, HIV LTR promoter transactivation, secretion of exRT, and expression of CA-HIV RNA species (Tat-Rev, Gag-Pol) mRNA in the HeLa-derived cervical cancer cell line, known as TZM-bl cells. These observations are interesting because HIV replication begins with the expression of short multiply spliced (MS) mRNAs encoding the viral regulatory proteins Tat, Rev, and Nef. Although HIV Tat is one of the initial transcripts made during HIV replication, as infection proceeds, HIV singly spliced (SS) mRNAs (encoding Env and the HIV accessory genes Vif, Vpr, and Vpu), as well as full-length unspliced (US) transcripts are produced. The unspliced HIV RNA serves as the mRNA encoding the Gag-Pol polyprotein and it is the virion genomic RNA. It is interesting that while 100 µg of BBR significantly inhibits both Tat-Rev mRNA which represents MS and Gag-Pol mRNA (genomic and Gag-coding), which represents US RNA species, 40 µg of BBR increased the levels of MS RNA (**Figure 6J, top**) but decreased US RNA (**Figure 6J, bottom**). Noteworthy, MS RNA, while it is the earliest transcript made, it may not measure the transcription and replication competence of HIV provirus [66]. The observed decrease in Tat mRNA level in response to treatment with 100 µg BBR was not due to decrease in cellular toxicity, because 100 µg BBR decreased HIV Tat and Gag-Pol to ∼46% and 6% normalized to GAPDH (used as endogenous control). These data imply that BBR may decrease the amount of genomic viral mRNA required for particle assembly, thus decreasing the production of infectious viral particles that can initiate the next round of replication. It is yet to be determined whether proviruses produced in the presence of 40 µg of BBR are replication competent.

We further showed that BBR inhibits Tat-mediated HIV LTR promoter transactivation, suggesting that in addition to the inhibition of the expression of CA-HIV RNA species, the mechanisms of BBR-mediated inhibition of HIV may also involve targeting HIV Tat-mediated HIV LTR promoter transactivation. The reduction of Tat-mediated HIV LTR promoter transactivation in the presence of exogenous rTat and HIV Tat mRNA suppression by BBR clearly suggests inhibitory role of BBR on Tat function. This observation is significant because Tat protein is not only critical for transactivation of HIV LTR promoter, HIV DNA transcription, and survival of infected cells, but mediates various cellular non-viral replication and non-AIDS-defining diseases (NADs) including cancer [7, 67]. Interesting, BBR showed concentration dependent inhibition of Tat-mediated cervical cancer cell migration and matrix invasion (**Figure 7**).

In our study, we showed that TZM-bl cells collectively migrate towards a mechanical wound area and HIV infection promotes the migration ability of TZM-bl cells. However, treatment with 100 µg of BBR significantly decreased cell migration resulting in decreased rate of wound closure in both infected and uninfected cells (**Figures 7D, 7E**). Similarly, the HIV Tat protein promotes cell migration and BBR inhibits Tat-induced cell migration. Interestingly, we also found that Tat promotes matrix invasion of TZM-bl cells and the cell clustering of the cells during invasion while BBR inhibits cell invasion and clustering as shown in invaded cells on the membrane (**Figure 7F**) and in the basal chamber (**Figures 7H, 7I**). Since collective cell migration and invasion depends on cell to cell interactions, BBR may inhibit collective cell to cell behaviors that depend on stable cell to cell linkages that promote pathogenic signaling. It remains to be determined the molecular variability of cell clusters, collective cell migration, and invasion, their different linkages to the cytoskeleton, and how BBR affects these behaviors and properties.

Insight into the potential mechanisms of BBR mediated HIV suppression and inhibition of Tat functions was provided by molecular docking and dynamic stimulation showing that BBR establishes π-amide stacking with Tat LYS71. It is noteworthy that π-amide stacking is as strong as hydrogen bonds and may play a critical role in protein–ligand binding. This observation is significant because LYS71 is a highly conserved residue in HIV Tat. LYS71is found in ∼74% of HIV-1 isolates across all HIV clades as reported in the HIV-1 sequence compendium [68]. Tat autoacetylates itself on LYS41 and LYS71 [69], the absence of which may reduce HIV replication by preventing the enhancement of histone acetyltransferase activity by p300 [69]. Moreover, polyubiquitylation at LYS71 is required for efficient HIV LTR promoter transactivation with no effect on the stability of Tat [70]. Mutating both LYS51 and LYS71 was shown to obliterate HIV LTR promoter transactivation completely [71]. Hence, the ability of BBR to control the activity of Tat is remarkable because HIV Tat protein is critical for efficient HIV transcription, latency, and rebound in PWH whose therapy was interrupted. Thus, any strategy that blocks HIV Tat may hold clinical promise in HIV cure.

## 5. CONCLUSIONS

In summary, our finding that BBR correlatively and significantly inhibits cervical cancer and HIV processes that may play important roles in HIV and cervical cancer progression is intriguing. This is because the prevalence of HPV-associated cancers, especially cervical cancer is high among women with HIV and the treatment of this cancer is limited to chemo and radiation therapies, lacks effective second- or third-line treatments, and not curative. Thus, compounds like BBR with the ability to inhibit both diseases may be developed to provide an alternative novel and effective treatment options. In future studies, we will assess the efficacy of BBR to inhibit HIV infection in primary cells that are targets of HIV, such human peripheral blood mononuclear cells (PBMCs), CD4+ T cells, macrophages, microglia, and monocytes. We will also use our unique resources, including genetically engineered mouse (GEM) models, patient derived xenografts (PDXs), and MmuPV1 mouse papillomavirus that models the pathogenesis and molecular activities high-risk HPVs [72] to test in vivo efficacy of BBR on cervical cancer and HIV comorbidity.

## Supporting information

Table S1 Stattistics for Tat-mediated migration

## Funding

National Institutes of Health grant R01DA042348-01 (to CMO)

National Institutes of Health grant R01DA050169 (to CMO)

National Institutes of Health grant R21/R33DA053643 (to CMO)

Startup funds from NYMC and Lovelace Biomedical

## Author contributions

Conceptualization: XL, CMO

Methodology: WN, BCO, HKI, ZZW, NY, XL, CMO

Investigation: WN, BCO, HKI, ZZW, NY, XL, CMO

Visualization: WN, ZZW, NY

Funding acquisition: XL, CMO

Project administration: HKI, XL, CMO

Supervision: HKI, XL, CMO

Writing – original draft: WN, BCO, HKI, ZZW, NY, XL, CMO

Writing – review & editing: WN, BCO, HKI, ZZW, NY, XL, CMO

## Competing interests

Authors declare that they have no competing interests.

## Data and materials availability

All data are available in the main text or the supplementary materials.

## SUPPORTING INFORMATION

**Table S1:** Statistics for Tat-mediated migration

## Notes

### Competing Interest Statement

The authors have declared no competing interest.

